# Existence of Causation without Correlation in Transcriptional Networks

**DOI:** 10.64898/2026.02.09.704821

**Authors:** Gerald M Pao, Ethan R Deyle, Hao Ye, Junko Ogawa, Marisela Guaderrama, Manching Ku, Tom Lorimer, Nina Tonnu, Erik Saberski, Joseph Park, Eugene Ke, Curt Wittenberg, Inder M Verma, George Sugihara

**Affiliations:** Okinawa Institute of Science and Technology Graduate University, Biological Nonlinear Dynamics Data Science Unit, Okinawa 904-0495 Japan; Scripps Institution of Oceanography, University of California San Diego, La Jolla, CA 92093-0202; Tufts University School of Arts and Sciences, Biology Medford, MA 02155; University of Pennsylvania, Philadelphia, PA 19104; University of California San Diego, Department of Pediatrics, La Jolla, CA 92093; Mammoth Biosciences, 1000 Marina Blvd, Brisbane, CA 94005; Department of Cell and Molecular Biology, The Scripps Research Institute, La Jolla, California, 92037; Salk Institute for Biological studies, La Jolla, California 92037; 9Department of Microbiology, Immunology, and Cancer, School of Medicine, University of Virginia, Charlottesville, VA 22903

**Author notes:** Correspondence: Gerald M. Pao, Okinawa Institute of Science and Technology Graduate University, Biological Nonlinear Dynamics Data Science Unit, 1919-1 Tancha, Onna-son, Kunigami-gun, Okinawa 904-0495 Japan, George Sugihara, Climate, Atmospheric Sciences, and Physical Oceanography Division, Scripps Institution of Oceanography, University of California San Diego, 9500 Gilman Dr., La Jolla, CA 92093-0202.

## Abstract

It is commonly assumed that lack of correlation is evidence for lack of causal relationship. Here, however we show that in transcriptional networks, causal linkages can exist in the absence of correlation. We find that a substantial proportion of transcribed genes in yeast and in mouse, show evidence of state-dependent (nonlinear) and temporally coordinated dynamics in their expression patterns (65-77%). Using a test that accommodates this fact, we uncover strong causal relationships that are invisible to correlation-based analyses for both yeast and mouse models. Specifically, for yeast we detect uncorrelated causal relationships for the transcriptional regulators *WHI5* and *YHP1,* and can verify these relationships experimentally. These genes reside at important checkpoints in the cell cycle where multiple signals are integrated at single nodes, giving rise to causal relationships, that despite being uncorrelated, can be accurately detected (71-78%) using a nonlinear causality test.

---

In the biological sciences and elsewhere it is often assumed that although correlation might not imply causation, correlation is absolutely *necessary* for causation – a view that finds special resonance when the relationships under study are fixed and unchanging or when the data are necessarily a-temporal. Despite its intuitive appeal, however, in many areas of natural science the robustness of this assumption as an operative guide is coming into question as time series data become more prevalent and complementary dynamic approaches to understanding causal relationships emerge. In a step toward assembling what we believe represents a more holistic view of how genes interact, we use genome-wide time series data of expression levels to uncover and experimentally verify causal links between *uncorrelated* genes that are thus invisible to correlation-based analyses and might otherwise go undetected.

Historically, time series of gene expression have been difficult and expensive to acquire. Molecular measurements typically destroy the sample, bulk populations of cells are difficult to synchronize [1], and until recently repeated large-scale measurements have been prohibitively expensive. Consequently, the vast majority of genome-scale studies have been cross-sectional, necessarily focusing on patterns emerging from static snapshots [2]. While more recent advances in both experimental techniques and data analysis suggest that longitudinal studies of time series could become increasingly prevalent (see Discussion), the historical contingency remains, with static cross-sectional data inviting a-temporal statistical analyses grounded in linear correlations. Such classical investigative protocols rely on the common-sense assumption that co-expressed genes are co-regulated [3–5]; meaning, for the most part, our main tool to explore the gene network has been statistical correlation.

However, there is evidence to suggest that even in such well-studied model systems as yeast [6], *S. cerevisiae* [7–10], as well as in mammalian immune cells [11, 12] and cancer cells [13], the process of gene expression goes beyond the standard a-temporal statistical domain and involves coordinated timing-, sequence-, and state-dependence. Thus, to understand the gene expression network in this coordinated temporal sense, we must account for how the activity of one gene affects the evolving expression of another, and more broadly how such ephemeral interactions are embedded in a wider network of changing cellular and extracellular influences. Importantly, there is increasing awareness that the transitory nature of these changing associations among genes and their interactions means that our common-sense fixed-time assumptions about causation and correlation (particularly that causes must be correlated with their effects) need not hold [14]. For example, as we will demonstrate below, it is possible that depending on context or temporal sequence, a positive association between genes may flip to negative, producing an overall lack of correlation. Although the general conditions supporting this phenomenon appear to be novel, there are indeed well-documented individual cases of genes that show a lack of correlation, despite being functionally related [12, 15]. Nonetheless, understanding the prevalence of this phenomenon and how it occurs remains largely unexplored.

To make these ideas concrete and to illuminate this potentially broad and consequential issue, we will examine data from both yeast (*S. cerevisiae*), and mammalian cells (mouse fibroblasts) specifically acquired to produce long time series of genome-wide expression. Acquiring these time series is accomplished here by using synchronized populations of cells that are repeatedly subsampled for RNA sequencing [16, 17]. Our general aim is to generate expression time series that closely approximate the genome-wide expression in the average representative cell (and do so without needing to account for dropouts of low expressed transcripts [18]). As described in Methods, for *S. cerevisiae*, cells are synchronized with alpha factor arrest and sampled every 5 minutes over 2 entire cell cycles, to produce data series having a total of 50-58 timepoints (see Fig. S1 for validation). For the mouse embryo fibroblasts, cells are synchronized through serum starvation, stimulation, and release, and sampled every 30 minutes over 3 cycles of arrest/stimulation, giving time series having a total of 122 timepoints. As a practical matter we place a particular emphasis in the yeast on well-studied genes sitting at cell cycle checkpoints as these were better synchronized and provide clean time series. Beyond their experimental convenience such genes are particularly interesting in that this is where signal-integrating behavior and thus coordinated control is likely to be important.

## Concrete insights from dynamic data

Armed with time series data on expression levels, it is now possible to better visualize how such time series can reveal relationships between uncorrelated genes. To begin, consider the case of the G1 cyclin CLN3 and the transcription factor SWI4 in our yeast data, both of which are important players at the G1 to S checkpoint [19]. Although there is no significant linear correlation between these genes in our data (Pearson ρ < 0.1, Fig. 1A), when the measurements are connected according to temporal sequence, an orderly relationship emerges (Fig. 1B). As suggested above, sometimes the variables are positively associated (at high levels of CLN3 and SWI4) and sometimes they are negatively associated (at low levels of CLN3 and SWI4); and this association varies sequentially through the cell cycle. This state- and sequence-dependence in the association between CLN3 and SWI4 expression is the hallmark of so-called *nonlinearity,* and as we see, nonlinearity can destroy linear correlation between dynamically interacting genes. One might hope that this is an isolated case, but as we discover below, nonlinearity is ubiquitous in our data.

**Figure 1.**
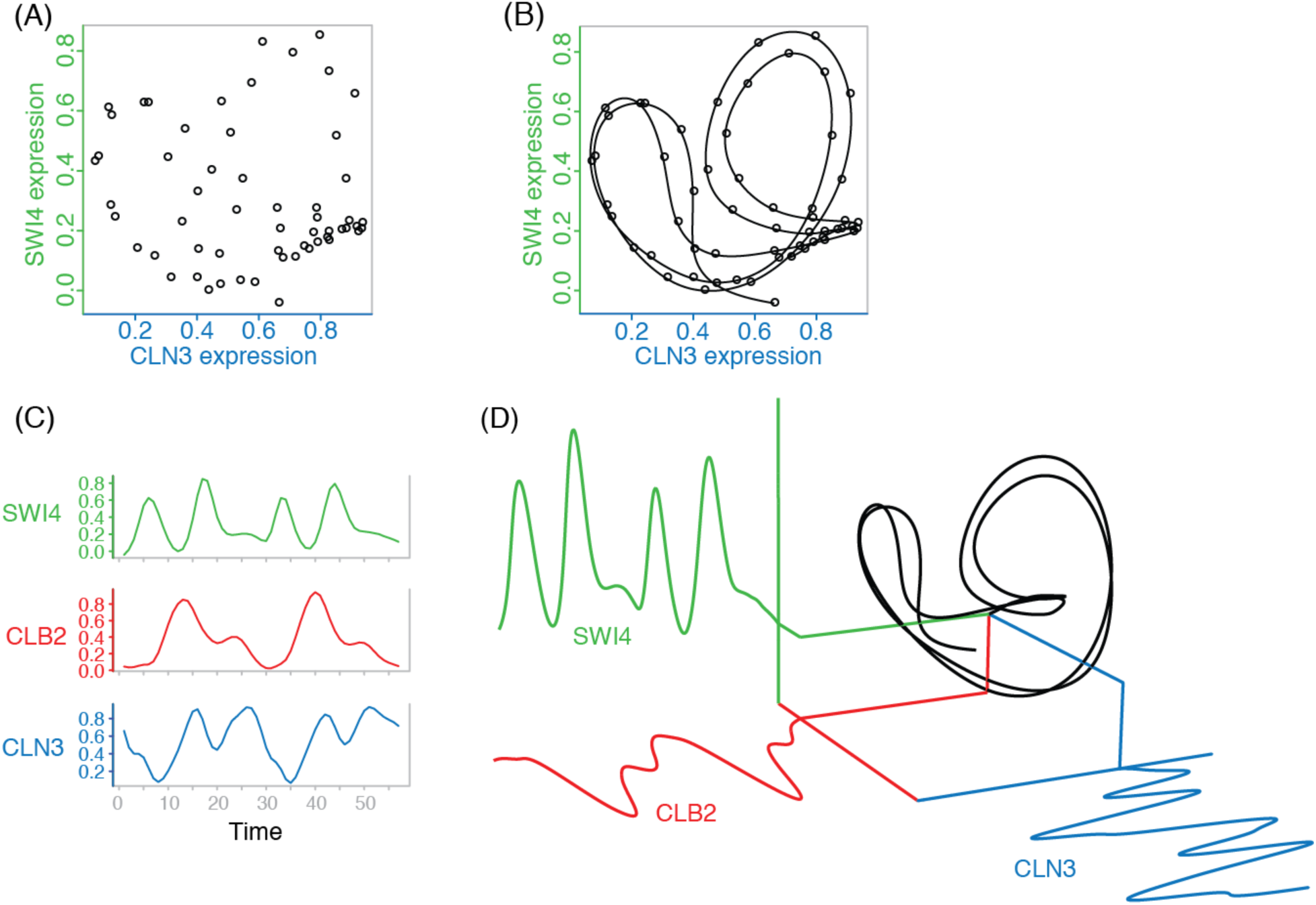
A dynamical representation of gene expression. (A) When viewed a-temporally, the normalized expression levels of transcription factor SWI4 and cyclin CLN3 are uncorrelated (ρ = - 0.047, n = 57 on the raw data). (B) However, connecting consecutive observations in time reveals a coherent trajectory describing a state-dependent (nonlinear) dynamic relationship between SWI4 and CLN3. Notice that there are points where the trajectory crosses itself and the future path is indeterminate (so-called singularities). (C) Expression time series for SWI4, CLN3, and the G3 cyclin CLB2, (D) Adding CLB2 as a third dimension that is known to interact with SWI4 and CLN3 untangles the points in (B) where the trajectory crosses itself. The resulting 3-dimensional attractor shows how the associations between genes (positive or negative) depend on location (system state). In general, individual gene expression time series can be understood as one-dimensional projections of multigene attractor dynamics onto a single coordinate axes. Note that unless otherwise stated, for visual clarity figures are drawn from temporally smoothed data then interpolated (see methods), though all analyses (numbers) are performed on un-smoothed data. (see supplemental_movie1 for raw data)

Although nonlinearity may appear to be a complicating feature that hinders understanding, it can in fact be an asset to be exploited in order to improve understanding [20]. Note that the trajectories in Fig. 1B reveal a dynamic relationship between CLN3 and SWI4 that is nearly deterministic. That is to say, knowing the expression level of both genes allows us to move forward in time along a trajectory that more or less uniquely predicts their near-term future expression levels. However, also note that when we do so there are places in Fig. 1B where the trajectories cross. These are points along the historical trajectory with non-unique futures where it appears the system can diverge non-deterministically and travel along either of the paths at the intersection. At such points, referred to as *singularities*, additional information would be required to determine which path the system will take. If the dynamics are deterministic this suggests that additional factors or dimensions are required to fully understand the system, as additional coordinates are required to resolve singularities. In this particular case, we know from previous studies that CLB2 interacts with CLN3 and SWI4 [21], thus when we include CLB2 expression as a third active dimension, the trajectory untangles (Fig. 1C). This untangled trajectory reveals with better clarity an underlying coherent geometric object – an attractor – that embodies the deterministic rules of the system. Just as with a system of coupled differential equations, an attractor reconstructed from data shows how the relationships between the three variables vary in time with changing cell states (see video clip https://youtu.be/fevurdpiRYg).

However, to reconstruct the attractor we first need to find an appropriate set of variables that identify the system – a set of *embedding* coordinates as in Fig. 1C (also referred to as active variables or native coordinates). The active variables in Fig. 1C were known in advance, but we usually don’t know which sets of genes interact or even have an idea of how many genes may be involved (this is in fact, largely what we are trying to discover). Fortunately, an answer to this problem can be found in a powerful analysis technique from dynamical systems theory. According to Takens theorem [22], information about the full dynamical system (i.e., set of interacting genes) is generically recoverable from a time series of *any one* of the coupled variables (i.e., observations on a single gene). Thus, in principle, a single gene time series can be used to construct a topologically similar attractor or “shadow” version of the original multi-gene attractor where time lags are used as proxy coordinates for the original variables (see video clip https://youtu.be/QQwtrWBwxQg and Fig S2). The shadow attractors are said to be diffeomorphic in that they preserve topological invariants such as the Lyapunov spectrum.

This process, illustrated in the video, can be applied directly to our data. For example, in Fig. 2A, shadow versions of the original attractor in Fig. 1C can be created by substituting lagged copies of the individual time series in place of the other native coordinates (e.g., native coordinates CLB2 and CLN3 can be replaced by lagged copies of the SWI4 time series, or lagged copies of CLN3 for SWI4 and CLB2). Each of these alternative univariate (single gene) reconstructions can be seen to be grossly similar to the original multiple gene native attractor, but more importantly each one provides a different (distorted but diffeomorphic) view of the same underlying system [23]. Most importantly, because they capture both the geometry and dynamics of the system’s attractor, these single gene representations can be used to assess whether the system is predictable and whether its dynamics are nonlinear [24].

**Figure 2.**
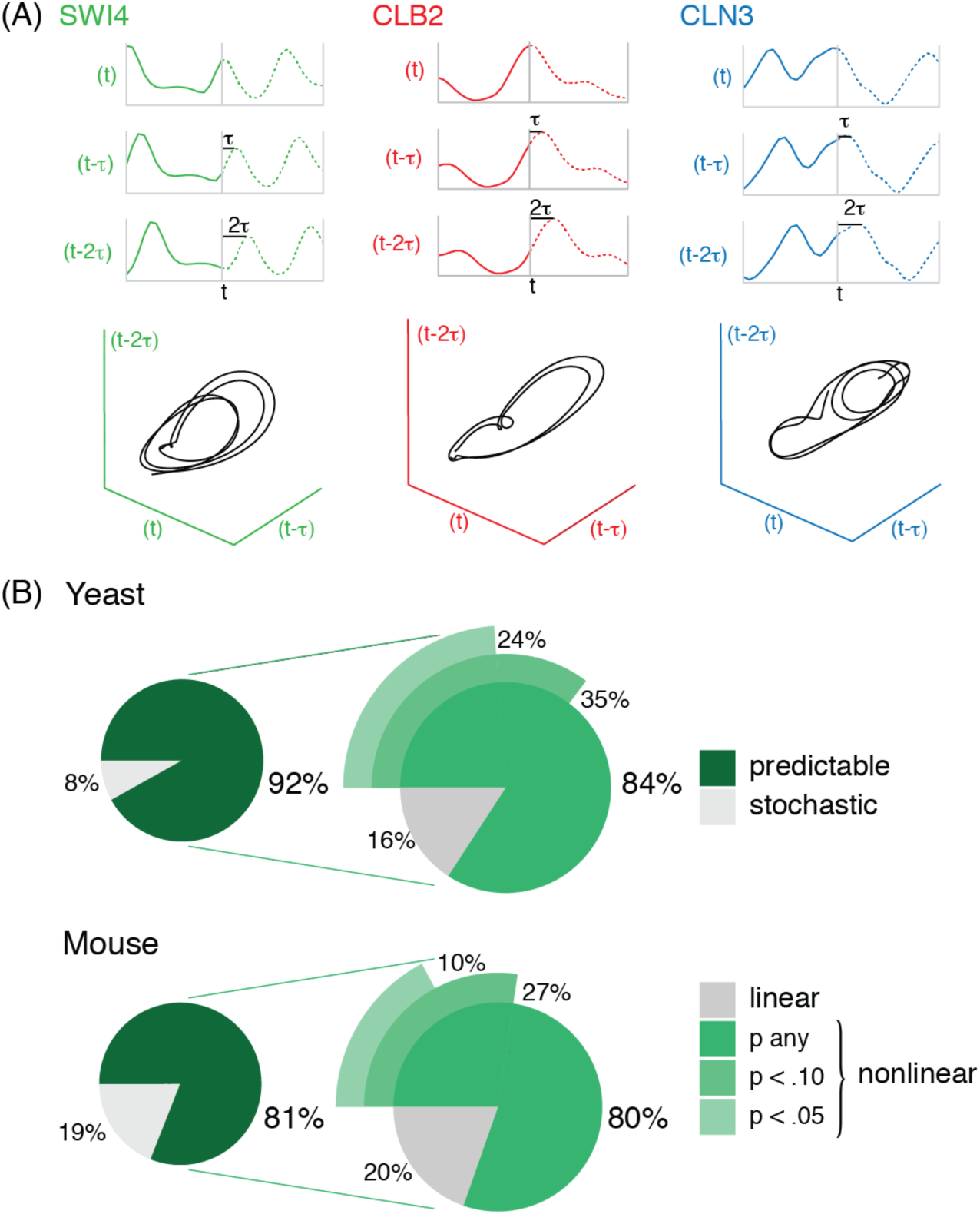
Testing the ubiquity of nonlinear dynamics in gene expression with single gene attractor reconstruction. (A) Three-dimensional lagged-coordinate (single gene) attractor reconstructions from smoothed expression time series for CLB2, SWI4, and CLN3 respectively. (B) The majority of collected time series in both yeast and mouse genes appear to be deterministically predictable, 92% and 81% respectively; and most of these genes have nonlinear dynamics, 84% and 80% respectively (see Methods). Details of these results are in Fig. S3.

## Prevalence of predictability and nonlinearity in gene expression

Fig. 2B summarizes the results of applying this single gene dynamic analysis to our yeast and mouse time series to obtain an empirical genome-wide assessment of the prevalence of nonlinear dynamics operating in these two systems. Nonlinear dynamics is detected in a time series if the reconstructed single-gene attractor is predictive (shows evidence of deterministic behavior) and if this predictability shows state-dependence (predictability that varies with the specific location on the attractor). We find that 92% of the 6,189 yeast genes and 81% of the 22,532 sampled mouse genes show significant predictability at the 5% level (see Methods). Applying an S-map test to assess the prevalence of nonlinear behavior in the significantly predictable genes (see Methods), we find that 84% of the yeast genes and 80% of the mouse genes show evidence of nonlinearity. It is important to note that these percentages represent conservative estimates as it is well known that the ability to detect predictability and nonlinearity in observational data increases with time-series length [25], and our time series are relatively short. The under-estimated prevalence of nonlinearity is apparent in our data (Fig. S3). Thus, far from being an oddity, the upturning picture of nonlinearity shown for CLB2, CLN3 and SWI4 expression in Figure 1 appears to be a ubiquitous genome-wide feature in these two systems. As such, nonlinearity and the fact that gene expression is dynamic sets the stage for how we study gene networks.

## Detecting causation without correlation in gene expression

In addition to assessing predictability and diagnosing nonlinearity, single-gene (lagged-coordinate) attractor reconstructions can also be used to confront the difficult challenge of identifying causal links in nonlinear systems [14]. It is clear in Fig. 3 that SWI4 and WHI5 lack statistical correlation (Pearson ρ < 0.1, Fig. 3A), and that their expression levels are not simply shifted in time (Fig. 3B). However, when these time series are viewed as products of a dynamical system (i.e., as observations of motion on an attractor), we find that the state of the SWI4 attractor (constructed univariately from a single gene) can be used to predict the specific contemporaneous value of WHI5 expression. Specifically, the nearest neighbors surrounding each state (point) on the SWI4 attractor, match up temporally with corresponding points on the WHI5 attractor that because of causal linkage are also nearest neighbors (Fig. 3C). In this way, it is possible to estimate the WHI5 state corresponding to any particular state of SWI4. Moreover, the accuracy of this prediction (Pearson correlation between true and predicted values) increases as more data are used construct the attractors (Fig. 3D), a property known as *convergence* that indicates a relationship beyond simple statistical association. If there is an underlying attractor, more data implies that nearest neighbors will be closer to each other (the density of points in the attractor increases), which allows for more accurate predictions. The ability to predict WHI5 from SWI4 shows that SWI4 has information about WHI5 – WHI5 left a causal imprint on SWI4. This conclusion is supported by a number of studies showing that SWI4 and WHI5 interact during the G1 phase with WHI5 acting as an upstream regulator repressing transcriptional activity in SWI4 [26, 27]. This method for identifying causal links by tracking time correspondence of nearest neighbors on attractors is called convergent cross mapping (CCM, see video https://www.youtube.com/watch?v=NrFdIz-D2yM) [14].

**Figure 3.**
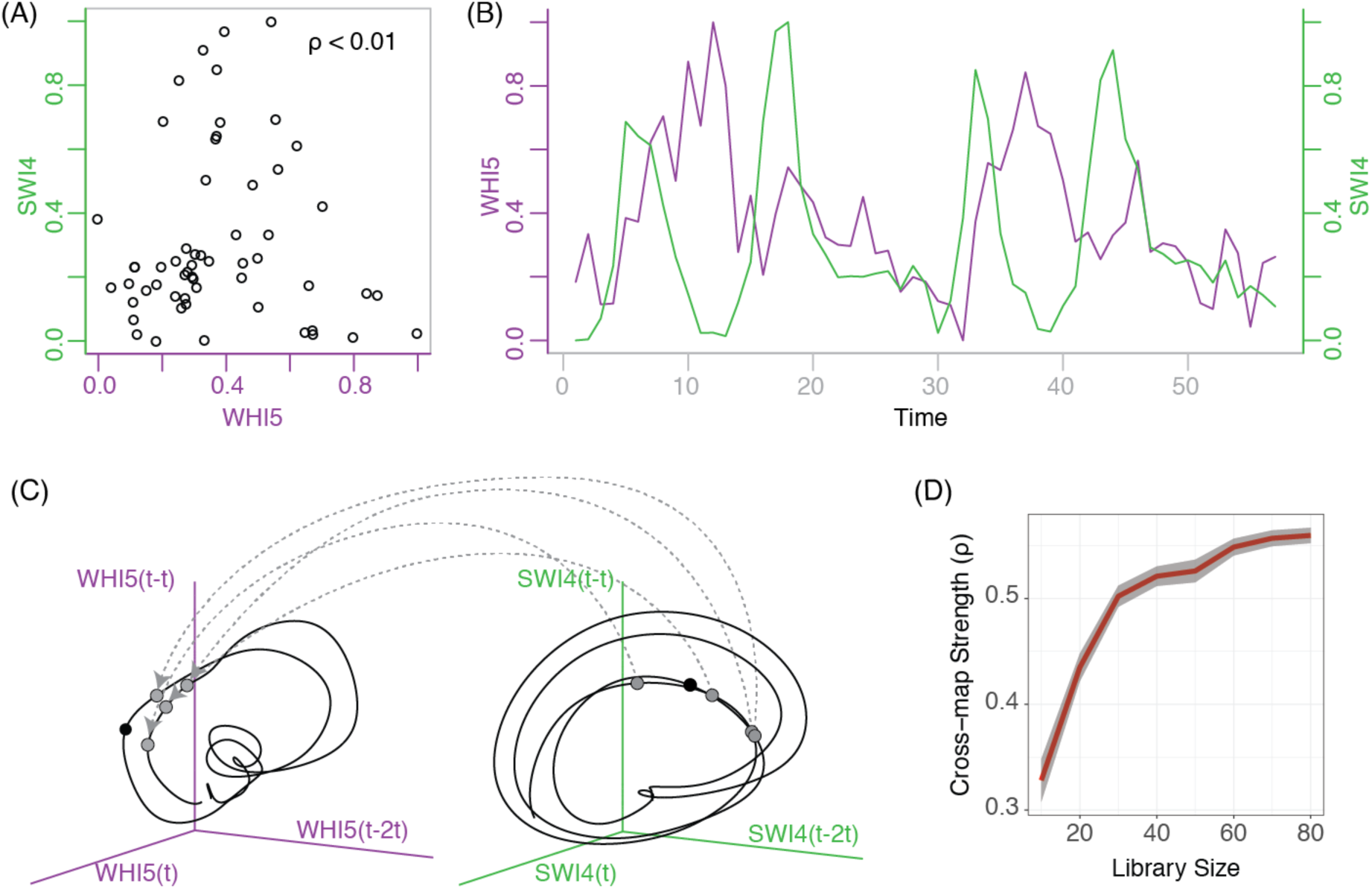
Uncovering WHI5’s causal effect on SWI4 by convergent cross mapping (CCM). (Note (A), (B) and (D) are pictured here with raw time series data). (A) WHI5 and SWI4 expression levels shows no discernable relationship (ρ < 0.1). (B) Lack of cross correlation between WHI5 and SWI4 is not simply due to a temporal shift. (C) Didactic illustration of cross mapping to test for a correspondence between single gene attractors. Specifically, if the nearest neighbors (light grey circles) of a particular illustrative time point (dark grey) on the SWI4 attractor correspond temporally to nearby neighbors of the same time point on the WHI5 attractor, then WHI5 expression values can be predicted from SWI4 values. This implies that WHI5 left a causal imprint on SWI4. (D) Result of application of this prediction process on raw wildtype data, showing cross map convergence (improvement in cross map prediction accuracy with increasing time series length). This verifies a true causal effect of WHI5 on SWI4, as opposed to a simple statistical association (where cross map strength would remain unchanged as a function of time series length).

More generally, when CCM is applied across all collected yeast and mouse genes that show evidence of nonlinearity, we find many cases where despite low correlation CCM will indicate the presence of significant nonlinear causal interaction (see Fig. S4 and Supplemental Dataset 1). That is, genome-wide application of CCM identifies many hidden but strong causal interactions between genes that appear to be robust. Fig. S5 shows that high CCM measurements are reproducible between experimental replicates of wildtype dynamics. While the robustness of CCM results is reassuring, a more satisfactory and definitive confirmation that CCM is accurate in identifying hidden causal links is to directly check them experimentally using genetic manipulation (over-expression or knockout).

## Experimental confirmation of detected causal links

We use two experimental approaches to determine whether manipulating an inferred uncorrelated upstream regulator provokes a response in inferred downstream targets. In the first experimental manipulation (ExM1), *WHI5* [26] was over expressed by placing it under control of the *GAL1* promoter (supplementary Fig. S1). For the second more direct experimental manipulation (ExM2), *YHP1* was simply eliminated in a knockout mutant. *WHI5* and *YHP1* were chosen based upon several criteria (see Fig. S1 caption), but most importantly because they are known to act at key checkpoints in the cell cycle where multiple signals must be integrated and this is where we expect causation without correlation can occur.

Time series of gene expression present an opportunity to examine the downstream effects of these manipulations in finer detail than would be possible with a-temporal data. Consider the yeast causal network, predicted by CCM, between WHI5 and the three genes examined earlier, CLN3, SWI4, and CLB2 (Fig. 4A). This network suggests that downstream effects of WHI5 overexpression should be visible in the time series of each of the other three genes, as indeed they are (Fig. 4B). However, in addition to apparent changes in mean level (e.g. CLN3), we also see subtler changes in timing and dynamics that are more difficult to quantify, and might not even be detectable with standard a-temporal expression data. When the timeseries for the three downstream variables are combined into the native attractor it is easy to see the dramatic change in structure before and after experimental manipulation (Fig. 4C). In order to quantify the change in gene dynamics resulting from experimental manipulation of an upstream gene (i.e. to examine individual causal links), we will focus on individual genes, and use single-gene (lagged-coordinate) attractors once again.

**Figure 4.**
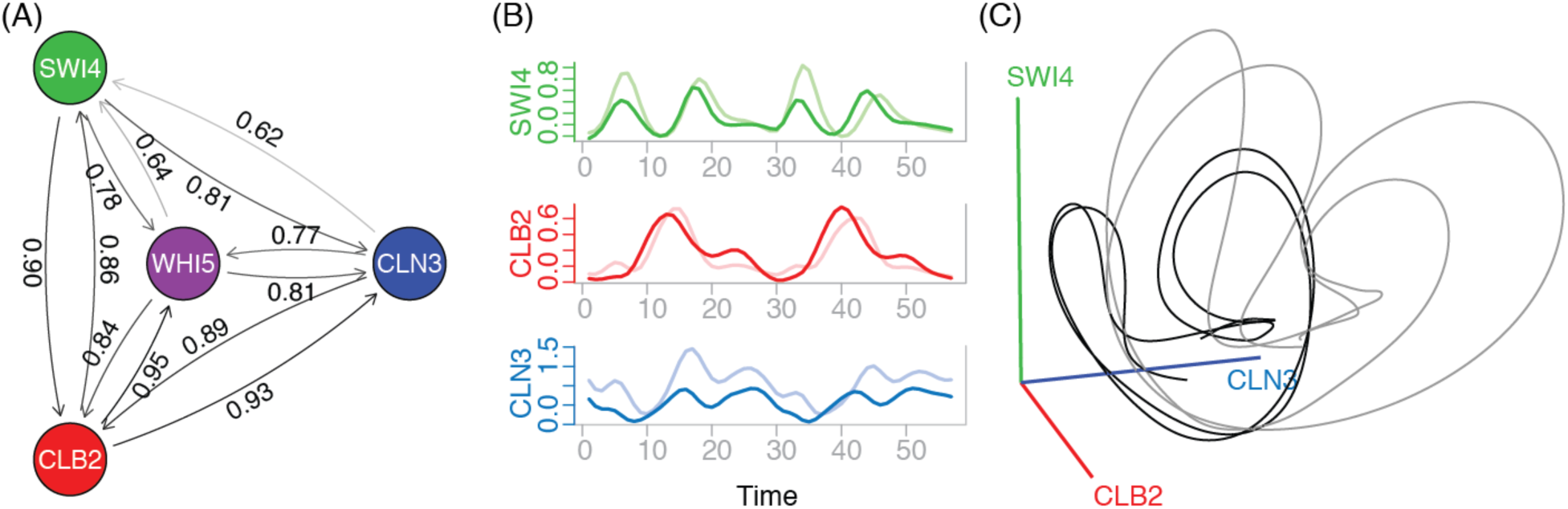
Confirming uncorrelated causal links with WHI5 overexpression. (A) A sub-network of transcriptional regulators (WHI5 and CLN3, SWI4, and CLB2) that are not mutually cross-correlated show bidirectional causal interactions. The strength of causal interactions is indicated by the Pearson correlation of cross map predictions made by CCM (numbers associated with arrows). (B) Changes in the dynamics of individual downstream genes CLN3, SWI4 and CLB2 from wildtype (dark lines) to WHI5 overexpression (light lines) as seen in their time series. (C) Combining the CLN3, SWI4 and CLB2 time series into a 3-dimensional attractor presents a clearer picture of how the wildtype attractor (black) deforms with WHI5 overexpression (gray) (see also supplemental_movie2 for raw data).

Focusing on the WHI5 to CLN3 causal link, Fig. 5A shows that the lagged-coordinate attractors generated from the two independent wildtype time series for CLN3 are indeed similar (Fig. 5A, left); however with experimental manipulation (WHI5 overexpression) the CLN3 attractor is vastly different (Fig. 5A, right).

**Figure 5.**
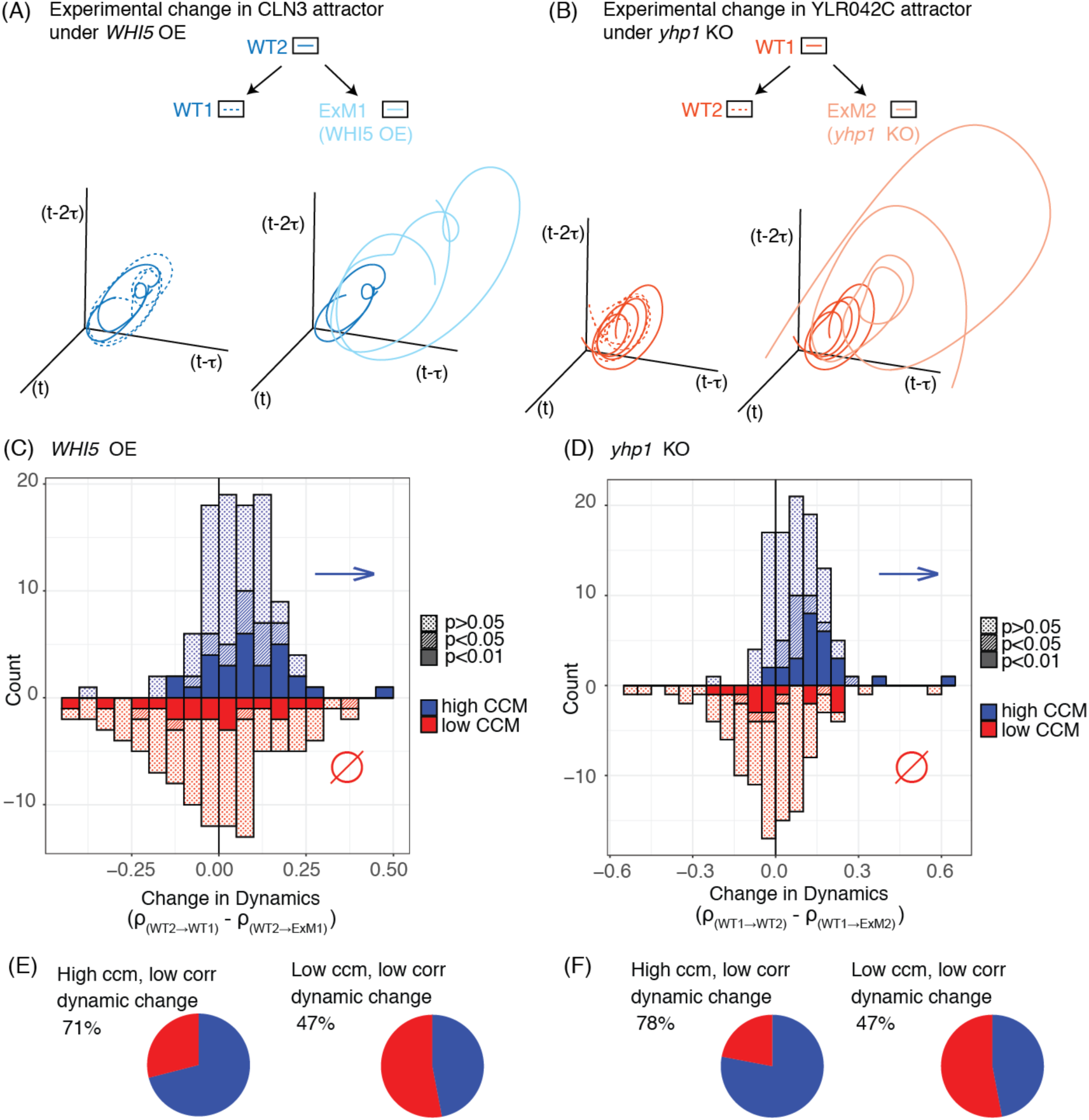
Statistical validation of causal links in two separate experiments: WHI5 overexpression (OE) and YHP1 knockout (KO). A predicted causal link is confirmed experimentally if there is a change in dynamics measured by a decrease in co-prediction skill between the wildtype and experimental time series (ρ*-diff* > 0, see text). (A) The co-prediction test is graphically illustrated for the predicted WHI5 target, CLN3. Here the wildtype attractor (WT2) is used as the predictor (reference library) to evaluate if the manipulated attractor (ExM1) is different from the wild type attractor from that experiment (WT1), the control. (B) Same as in (A) but where WT1 is the reference to evaluate the difference between the ExM2 and WT2 attractors for the YHP1 target YLR042C. Note that the striking differences between experimental manipulation attractors and wildtypes is not related to data smoothing (see Fig. S8). (C) and (D) Co-prediction under WHI5 OE (C) and YHP KO (D) are examined for the genes showing low correlation (ρ < 0.1). The distribution of ρ-diff for the 100 highest CCM genes (blue) shows a significant positive shift in both cases (p < 10^−8^ under Wilcoxon test). Conversely the low CCM genes do not show a significant shift of ρ-diff from zero. This indicates that the detection of causation without correlation by CCM is statistically significant. (E) and (F) summarize differences between the groups of genes.

With this setup, changes in dynamics due to experimental manipulation can be effectively measured by how well or poorly the manipulated time series can be predicted from an attractor (a predictive reference library) built from an unmanipulated wild type time series. This method, known as *co-prediction* [28, 29], quantifies dynamic similarity between attractors (see Methods). It differs critically from cross-prediction used in CCM in that the implicit hypothesis is that the two time series (the reference and the manipulated) are actually the same variable (different realizations of the same underlying attractor dynamics, up to a linear scaling factor), rather than interrelated or dynamically coupled variables. Thus, to test whether variables *X*_1_ and *X*_2_ are co-predictable one uses {*X*_1_} to predict {*X*_2_} and vice versa, possibly normalizing each time series first (see Data Processing Standardization in Methods). The skill of co-prediction, measured by the Pearson’s correlation between observed *X*_2_ to co-predicted *X*_1_, is a measure of the similarity of the dynamics of *X*_1_ and *X*_2_. Here, to further minimize possible dynamically sensitive batch effects such as one experiment producing slightly noisier outputs (e.g., lower predictability suggests that the knockout experiment was slightly noisier) we evaluate the experimental manipulation relative to the wildtype from the same experiment (the control). This requires using a common reference library, which we choose as the wildtype from the other experiment. Thus, to assess change under the Batch 1 experiment (WHI5 overexpression), WT1 is taken as the control, ExM1 is taken as the manipulation, and WT2 (the wildtype from the YHP1 knockout experiment) is used to construct a common reference attractor. That is, to assess WHI5 overexpression dynamics a reference attractor constructed from WT2 is used to co-predict both WT1 and ExM1 (see box and arrows diagram at the top of Fig. 5).

In Fig. 5A, left, where the dynamics of the independent wildtypes should in fact be the same, the co-prediction skill is high (ρ_WT2→WT1_ = 0.853). Conversely, in Fig. 5A, right, where the attractor from the wildtype dynamics differs substantially from the manipulated dynamics produced by WHI5 overexpression, the co-prediction skill is lower (ρ_WT2→ExM1_ = 0.653). A positive difference between these co-prediction skills, ρ_diff_ = ρ_WT2→WT1_ – ρ_WT2→ExM1_ indicates that WHI5 overexpression has changed the downstream dynamics of CLN3, thus verifying the existence of the causal link. Fig. 5B shows a similar example for YLR024C under YHP1 knockout, where ρ_WT1→WT2_ = 0.658 and ρ_WT1→ExM2_ = 0.353 resulting in a positive ρ_diff =_ ρ_WT1→WT2_ – ρ_WT1→ExM2_, which again verifies the inferred causal link.

To assess the statistical significance of the ability of CCM to correctly discriminate causal from non-causal linkages (again, where is no correlation (ρ < 0.1) with the manipulated genes), the analysis was repeated across genes that were either among the top 100 (strongest uncorrelated CCM genes) or bottom 100 (weakest uncorrelated CCM genes) with respect to causal linkage to each of the manipulated genes: WHI5 or YHP1. If CCM can successfully uncover hidden causality, then the genes where CCM reveals a strong causal connection to the manipulated gene should on average show positive ρ_diff_, whereas those for which CCM did not reveal significant causation should on average show zero ρ_diff_. This indeed is the case for both WHI5 overexpression (Fig. 5C) and YHP1 knockout (Fig. 5D), where the distribution of ρ_diff_ for the 100 highest CCM uncorrelated genes (blue) is substantially positively shifted (p < 10^−8^ under Wilcoxon test in both cases) and the ρ_diff_ distribution for the low CCM uncorrelated genes (red) is not significantly shifted from zero. Accordingly, under WHI5 overexpression and YHP1 knockout, 71% and 78% of the high CCM uncorrelated genes respectively show ρ_diff_ > 0, compared to 52% and 44% of the low CCM uncorrelated genes (Fig. 5E, F). Altogether, these results confirm the statistical significance of our ability to detect causation without correlation in the wildtype time series.

## Revisiting well-studied gene networks

The fact that causation between uncorrelated genes can be detected from time series data, invites revisiting other well-studied networks. Could this phenomenon also apply in a mammalian example? Figure 6 shows a CCM analysis on a well-studied group of genes related to the inflammatory response in our mouse embryonic fibroblast serum response time series. In this example, cross correlation yields a somewhat ambiguous network picture of the known regulatory relationships for the NF-kB system, in that RelA and IkBα are only weakly correlated even though it is known that there is negative feedback regulation by IkBα where NF-kB exhibits oscillations that decay over time [11, 30] (Fig. 6A). CCM on the other hand correctly identifies IkBα as the target of RelA and the feedback is seen as an asymmetric interaction with RelA having a very strong effect on IkBα but not vice versa (Fig. 6B). This is consistent with the fact that IkBα is known to be a direct transcriptional target of the RelA/p50 heterodimer transcription complex. Although IkBα is a strong inhibitor of RelA at the posttranscriptional level where IkBα protein sequesters RelA/p50 protein and maintains it in an inactive form in the cytoplasm, IkBα was not thought to affect RelA transcriptionally, but rather is thought to be constitutively expressed. These results suggest that IkBα expression may exert a weak albeit detectable regulatory influence on RelA that may very well be indirect.

**Figure 6.**
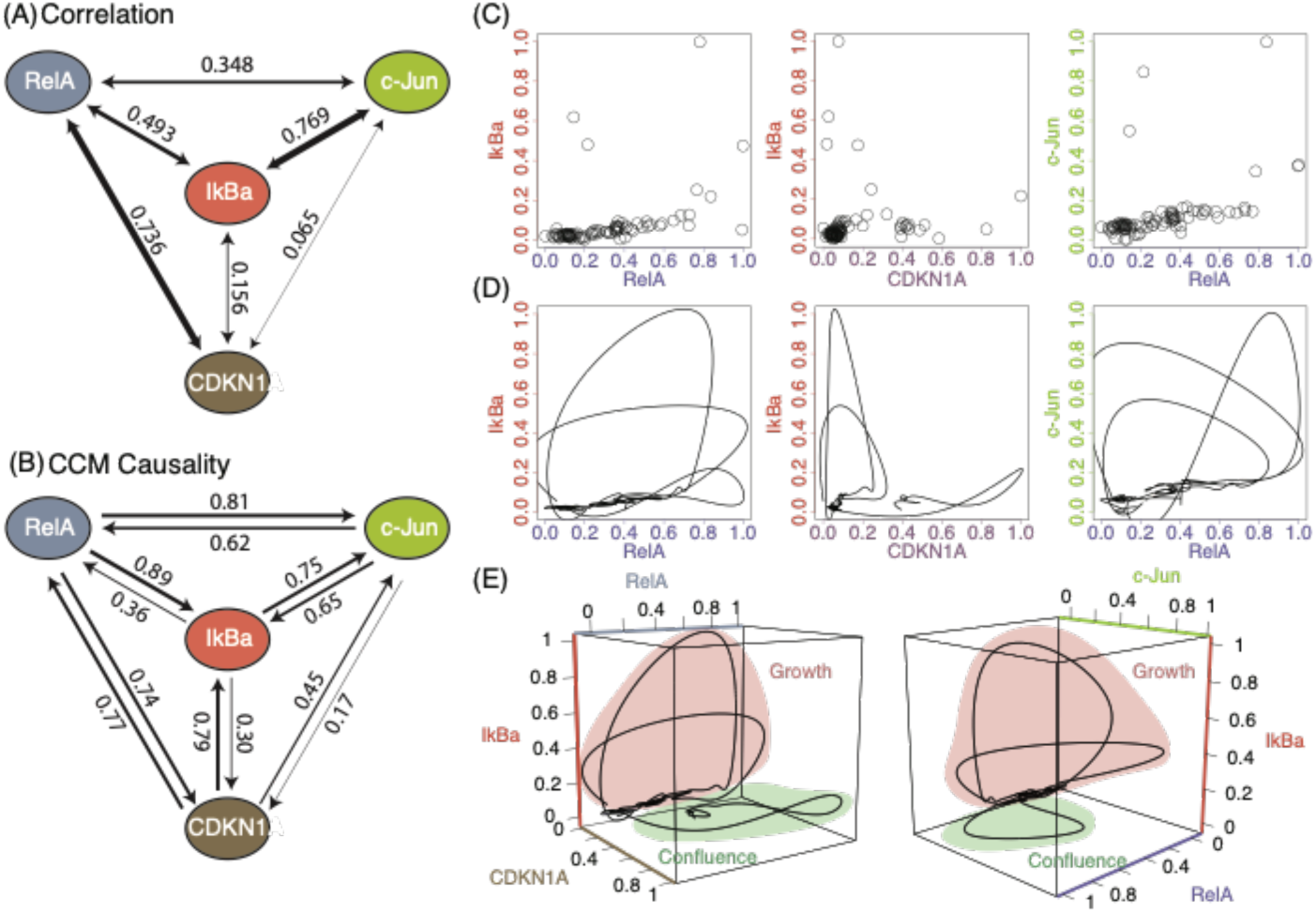
CCM uncovers novel causal links in the NFKB response network. (A) Network based on cross correlation between RelA, IkBα, c-Jun and CDKN1 (numbers associated with arrows show symmetric cross-correlation). Despite experimental evidence for the interaction between RelA and IkBα, they are only weakly correlated (ρ = 0.493). Weak correlation is also seen between IkBα and CDKN1A (ρ = 0.156) as well as c-Jun and RelA (ρ = 0.348). (C) Scatter plots of expression between these pairs shows weak associations, (D) but dynamic trajectories suggest structured relationships. (E) 3-dimensional attractor reconstructions show how different cell states occupy different parts of the attractor during exponential growth phase and confluence. CCM detects the known relationship between RelA and IkBα while also suggesting causal links between IkBα and CDK1NA and c-Jun and RelA.

Significantly, the CCM network identifies a previously undocumented link from RelA to c-Jun. Indeed, plotting RelA, IkBα and c-Jun together shows clear evidence of coherent attractor dynamics (Fig 6E). We also observe that the nonlinear relationships between the four genes RelA, IkBα, CDKN1A (P21) and c-Jun appear to show an additional source of state dependence: whether the cells are under exponential growth conditions or at confluence (Fig. 6E). Thus, even in an example as well-studied as the NF-kB system, by adopting an empirical dynamic perspective we are able to open up new avenues for investigation.

## Discussion

The time series data we analyzed indicate that a majority of changes in expression are neither random nor simply oscillations around a stable equilibrium, but appear to reflect nonlinear deterministic dynamics. Thus, even if firing of polymerase leading to gene transcription is stochastic [31, 32], changes in expression, assumed to be proxies for gene activity, are in fact predictable and indicative of the functional machinery dynamically coming together and falling apart over time. Such behavior should fundamentally change how gene expression is observed, analyzed, and understood. This dynamic view is complementary to the enormous experimental advances in genome-wide technologies that have led to an extensive and growing parts-list of the genetic units of the cell. Likewise, our methodology is distinct from but complementary to the normal deductive mode of perturbation and computer modeling that has defined systems biology [33], as it does not require a model having a particular functional form (models as hypothesis sensu [34]), nor does it require the rigorous determination of parameters necessary for a system of differential equations [34]. Rather, taking an inductive empirical dynamic approach we seek to address the vexing question: which of the tens-of-thousands of genes in the parts-list are causally related?

Other investigators have found compelling evidence in a-temporal expression data, for low-dimensional manifold geometry and nonlinear interactions [35], e.g. that are predictive of the expression levels of under-sampled genes [36]. The time-series approach taken here is a complementary perspective, with some key differences. First, the attractors here are built from the bottom up, from single genes found to be dynamically linked (mechanistically causal) rather than reduced from high dimensional data involving a large constellation of different genes (such data are often reduced to lower dimensional principal combinations of genes, to make them more interpretable and experimentally translatable). Second, instead of only obtaining geometry (the manifold representing a cloud of points) we explicitly obtain both geometry and dynamics. That is, we obtain the full attractor with the experimentally observed (not inferred) temporal vectors showing direction of flow. Finally, and as a consequence of the above, CCM allows us to address realized multidimensional *dynamics*, and by so doing can identify more complicated interactions that would otherwise be missed if attention was restricted to seeking simple one-dimensional linear (or nonlinear) relationships (e.g. Fig. 1A). Taken together, these differences allow us to address causality in a holistic context where the gene network is itself dynamic and resides within changing cellular and extracellular influences. Exploring how gene expression networks behave as dynamic (ephemeral) entities is a prime future avenue for investigation. Thus, beyond recovering causal networks for gene expression, a future focus will likely be on understanding how the specific linkages and thus the network itself can change [37].

As we have suggested, our estimates of the prevalence of nonlinearity are conservatively low. Aside from the previously discussed difficulty of detecting nonlinearity in short time series (see Fig. S3 [25]), our method for generating time series means that we are restricted to analyzing only those expression patterns that could be well synchronized in culture. For example, this restriction eliminates capturing the most nonlinear behavior (unstable with rapidly diverging future trajectories), as unstable behavior does not readily synchronize – behavior that is liable to be diluted or washed out by population averaging [38]. Conversely, and somewhat problematically, the most coherent signal to emerge with population averaging will tend to be tied to synchronized behavior, a fact that would artificially inflate the prevalence of linear correlations and diminish measurable nonlinearity. With so much stacked against detecting nonlinearity, the fact that these data show it to be ubiquitous speaks to how dominant it must be as a property of gene expression at large.

Our experimental validation of CCM is also conservative in that only one type of experimental manipulation was used to examine the predicted downstream effects for each examined gene (either overexpression or knockout). A single directional manipulation may not be sufficient to provoke a response in all causal targets. For example, if under normal circumstances WHI5 is not limiting to CLB2, then despite causal linkage, overexpression of WHI5 might have no effect, while knockout of WHI5 might show an effect.

Of particular interest is the fact that the overwhelming majority of uncorrelated but causal interactions identified by CCM (Supplemental Dataset 1), are cases where multiple drivers converge onto a single target (Fig. S7). This is potentially significant since the implied network structure is suggestive of signal integrators [39, 40]. Insofar as this connection is true, it is important to note that these putative nodes are highly nonlinear (Fig. 2), representing genes that exhibit state dependent expression patterns influenced by the coordinated expression of multiple upstream genes. This is illustrated in Fig. S7, where YHP1, a mediator of the M to G1 transition, takes in signals from 8 genes (that are all uncorrelated with YHP1), that in turn influence the expression of 3 downstream genes (that are also uncorrelated with YHP1). Such nonlinearity in the response of potential signal integrators has implications for identifying genes associated with check points and it may even be important for adaptation in the evolutionary context [41].

The ability to identify causation without correlation can enable the efficient discovery of genes related to cellular information nodes that are responsive to multiple inputs (and that are potentially signal integrators). This capability alleviates experimental burden and clearly shows an avenue forward by identifying which components need to be simultaneously manipulated to achieve a desired downstream response. For example, without guidance as to the underlying causal network, in order to evaluate all of the possible 3-gene combinations in a mammalian genome by brute force would require over 10 trillion experiments. Being able to narrow the search with an agnostic causality analysis such as this one represents a substantial step forward.

This gain in efficiency is all made possible by the relatively recent availability of time series data. The ability to generate this kind of data to study gene expression and map the functional genome is evolving quickly. While in this work we used destructive sampling of synchronized cell cultures to generate time series, there is also interest in generating pseudo-time series that approximate single-cell expression without requiring bulk sample synchronization. This is particularly applicable in cases where it has been possible to identify genes that vary slowly through cell differentiation which is essentially a one-way sequential expression cascade [35, 42–45]. There is also emerging work using light microcopy on live cells with fluorescent proteins to generate expression or activity time series for single cells [46]. Moving forward, as experimental technologies in molecular biology advance and more accurate time series data are collected, and such data become more common, analytical tools such as those employed here that address nonlinear dynamics in gene expression in a holistic and mechanistic way (bottom up rather than with statistical dimension reduction protocols) will become more informative and more essential to a deeper temporal understanding of gene expression in nature.

## Acknowledgements

This study was supported by the following sources: National Science Foundation (NSF) DEB-1655203, NSF-ABI-1667584, DoD-Strategic Environmental Research and Development Program (SERDP) 15 RC-2509; (GS, ED, HY) and the McQuown Chair in Natural Sciences, University of California, San Diego (GS); the Deutsche Forschungsgemeinschaft (SFB1160 “Immune-mediated pathology as a consequence of impaired immune reactions [IMPATH]”, project Z02 to M.K.). GMP, JO, NT and IMV were supported, in part, by a grant from the NIH (R01 CA095613); a Cancer Center Core grant (P30 CA014195-38); Ipsen; the H.N. and Frances C. Berger Foundation; the Leona M. and Harry B. Helmsley Charitable Trust grant #2017-PG-MED001, and MCK and the NGS Core Facility of the Salk Institute, with funding from NIH-NCI CCSG: P30 014195, and the Chapman Foundation. CW and MG were supported by grant (4R01GM059441-16) from NIH NIGMS.

## Supplementary Materials and Methods

### Protocol for Mating Pheromone synchrony

Cultures of the designated strain were grown to mid-logarithmic phase in YP-GAL (YEP with 2% galactose) medium and then treated with 50 ng/ml of alpha-factor for approximately 100 minutes or until cells were >95% unbudded. The cells were collected by centrifugation and resuspended in fresh YP-GAL medium lacking mating pheromone and sampled at 5-minute intervals by vacuum filtration and immediate chilling in ice cold water. Cell samples were collected by brief centrifugation and frozen in liquid nitrogen for subsequent preparation of RNA. An equivalent cell sample was taken for each time point and preserved with formaldehyde for counting. When cells achieved greater than 90% budding, mating pheromone was again added to 50 ng/ml and cells treated in the same manner as stated previously to resynchronize followed by release into fresh medium for the second round of division. Samples were collected through the next budding maximum until the proportion of budded cells decreased to less than 10%. Prepapration of samples for sequencing were performed in two experimental batches, each with its own WT control. Batch 1 consisted of 57 timepoints with CWY2090 and CWY2559 where WHI5 was overexpressed and batch 2 with CWY2090 and CWY2565, where the latter is a *yhp1Δ* mutant. Time points per two cycles ranged from 50-58 timepoints. Total RNA was isolated using RNAeasy (Qiagen) and polyA primed TruSeq libraries were constructed as described above. Individual RNAseq reactions were sequenced at a depth of 3 million reads/timepoint which allowed adequate quantification of S. cerevisiae gene expression for vast majority of all genes.

### Strain list

Wild Type: (CWY2090: *MATa his3Δ1,leu2Δ0,met15Δ0,ura3Δ0 bar1Δ::LEU2*)

*GAL-WHI5*: (CWY2559: *MATa his3Δ1,leu2Δ0,met15Δ0,ura3Δ0 bar1Δ::LEU2 GAL1-WHI5:URA3*)

*yhp1Δ*: (CWY2565*: MATa his3Δ1,leu2Δ0,met15Δ0,ura3Δ0 bar1Δ::LEU2 yhp1::KANr*)

### Mammalian serum response sample preparation

Mouse embryonic fibroblasts were generated as follows: Thirty female CF1 mice were mated and checked for plugs to time the pregnancy. Timed pregnant females were then sacrificed at day 13 of the pregnancy and embryos isolated. Mouse embryonic fibroblasts were then prepared by removing internal organs and decapitated and remaining carcasses minced mechanically before enzymatic treatment by collagenase and pronase to release the fibroblasts. Fibroblast were then plated and passaged once before frozen for storage in 10% DMSO and 90% serum in liquid nitrogen for long term storage. The fibroblasts were designated as passage one at the time of freezing. For the serum response experiment, we thawed fibroblasts and starved these for 72hours in 0.5% serum. These were subsequently trypsinized and plated in 120 30mm dishes for subsequent sampling. Cells were kept in 0.5% serum for a further 5 hours to insure attachment onto the plates. Initial samples were collected at this point to determine baseline gene expression levels. Samples were collected by aspiration of the growth medium, a single PBS wash and solubilization of the cells in Trizol scraping them off with a cell lifter and immediate freezing in dry ice. Cells were stimulated with 20% fetal calf serum for 24 hours before starving them again with 0.5% serum for 48 hours. Sampling was initially done at 0.5 hr intervals, but it was recognized this led to excessively high autocorrelation between time points. Thus, the final phase of sampling was done at 1 hr intervals for efficiency. Specifically, samples were collected at −1 hour to capture the state before stimulation, then at the first hour and then every 0.5 hours thereafter until 38 hours. The last 14 hours of these fall during the second period of serum starvation. The process was then repeated for a second cycle and samples were collected at −1 hr, then at time point zero and every 0.5 hours thereafter until 3 hours. Thereafter samples were collected every hour to 27 hours to complete a total of 96 samples collected. To test the serum response after confluency an additional 25 timepoints were collected at 0.5 hour intervals. Cells were left confluent for 72 hours under serum starvation conditions 0.5% serum and stimulated with 20% serum to initiate serum response. Samples were collected from the 0 timepoint before stimulation for up to 14 hours with 30-minute intervals. Total RNA of the samples were then purified using the Direct-Zol 96 well RNA isolation kit (Zymo Research) as described by the manufacturer. Amounts of RNA recovered were in all cases in excess of 1 microgram for subsequent processing. RNA quality was determined by Agilent TapeStation and all samples have RNA integrity numbers (RIN) of 8 or higher. Total RNA was used for the generation of polyA-selected mRNA RNAseq libraries using the Illumina TruSeq stranded library prep kit. Library was quality controlled by TapeStation to confirm the absence of primer dimers. All 96 samples were pooled and sequenced on 8 lanes of an Illumina HiSeq2500 sequencer at single end 50 base pairs with a target depth of 30 million reads per sample. Sequencing reads were then mapped onto the mouse genome (mm9 or mm10) and reads were quantified as transcripts per million reads (TPM). TPM values were then used for the generation of time-series data. For the EDM and causal analyses, we dropped half-hour timepoints from the higher frequency portions of the time series due to excessively high autocorrelation at 0.5 hr lag. Consequently, the time series analyzed consisted of 69 time points out of the initial 96 observations.

### Data Processing

#### Standardization

The raw RNAseq time series are first processed to account for potential batch effects, while preserving any changes in levels or dynamics between corresponding wildtype (WT_k_) and manipulated experiments (ExM_k_).

For gene *Y* of batch *k*, we apply a linear scaling to all time-points of *Y*_WTk_ and *Y*_ExMk_ so that the range for the scaled wildtype time series *Ỹ*_WTk_ is [0, 1]:

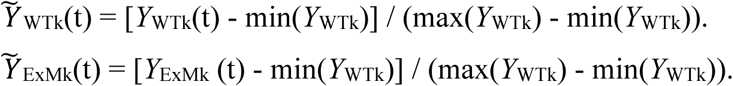

All remaining analyses are done on these transformed values *Ỹ*.

#### Selection

The standardized RNAseq time series were also filtered to eliminate time series with very poor resolution or measurement. A total of 818 time series having either near-zero variance (σ^2^ < 0.00001) or over half zeros were deemed as having essentially no signal and were discarded.

#### Smoothing and interpolation for visual presentation

Cubic splines were used to smooth and interpolate individual gene time for visual presentation in main text figures. The data observations are smoothed first with the function ‘smooth.spline()’ from the R ‘stats’ package 3.6.0 with smooth parameter *spar* = 0.4. All other parameters are left at default values. The purpose of this transformation is solely to reduce the effects of observational error on the visual presentation of individual gene trajectories. After observation points were smoothed, cubic interpolation is done to generate 20 points between observation intervals using ‘splinefun()’ and ‘spline()’ in the R ‘stats’ package 3.6.0 to visually approximate the trajectory of the gene dynamics in continuous time. All parameters are left at default values. Though these transformations can aid visual presentation of data, spline smoothing and interpolation is not used for any quantitative analysis here and is not generally recommended for EDM analysis, particularly where nonlinearity is being assessed, as such smoothing can introduce artificial linearity to the output.

### Analysis Calculations

#### A brief explanation of attractor reconstruction

Complex systems are often modeled using differential (or difference) equations that describe the transition through time between different states of the system. Each *<u>state</u>* is represented as a vector of state variables <u>x</u>(*t*). In ecology this might be abundances of foxes, rabbits and grasses etc. while for molecular systems this might be RNA and protein concentrations expressed by different interacting genes [53]. Regardless, the set of all states that a dynamic system transitions through forms a geometric construct known as an attractor *<u>manifold</u>*, ***M***. The attractor describes how state variables relate to each other through time. If there are rules governing how the system changes (if the genome is a dynamic system that is not *purely* random) there is an attractor to be uncovered (Figure S2A) [23]. Attractor manifolds determine (express) relationships among variables, and can be obtained simply by re-plotting the time series data (see video clip https://youtu.be/fevurdpiRYg). Constructing attractors empirically from gene expression time series is the basis of the empirical dynamic approach in this work.

In this attractor-based view of complex systems, a time series {*Y_i_*} is a projection of the dynamics occurring on ***M*** (Fig. S2B). More generally, the *Y_i_* are *<u>observation functions</u>* of the dynamics on ***M***. The *Y_i_* may be fundamental coordinates or they may be lagged coordinates or any function (e.g. rotations or linear combinations of the original Cartesian coordinates) that maps points in ***M*** to time series observations. The key insight is that biological time series can appear complex because they are projections into one dimension of dynamics occurring in higher dimensions.

If all the variables and equations governing an system are known we could construct the attractor manifold by direct simulation. In fact, it is also possible to reconstruct the manifold empirically, if we have time series for all the variables. This manifold would be an empirical expression of all of the dynamic relationships among variables observed in the data. However, in practice we may only have time series information about one species. A key result from dynamical system theory — the Takens embedding theorem — proves that one can reconstruct the dynamical attractor for a system from data in the form of time-lagged samples of just one variable, such as *Y_i_*. Thus, *<u>state space reconstruction (SSR)</u>* is a method to recover an approximation of ***M*** from time series. This is illustrated in the video clip https://youtu.be/QQwtrWBwxQg and in panel c of Fig. S2, where the shadow manifold, ***M***_1_’ is constructed using lags of time series {*Y*_1_}. The reconstruction captures the essential topology and dynamics of the original system. Once the attractor is reconstructed, simple non-parametric methods can be used to learn the dynamics, such as nearest-neighbor prediction [47], locally weighted linear regression [24], radial basis functions [48] and Gaussian processes [49]. This combination of attractor reconstruction with non-parametric modeling is referred to as Empirical Dynamic Modeling.

#### Univariate analysis

The presence of low-dimensional, nonlinear attractor dynamics in yeast and mouse MEF is quantitatively tested with a basic two step procedure [50]. In the first step, the simplex method [47] is used to assess predictability, and estimate the optimal embedding dimension. In the second step, the S-map method [24] is used to assess nonlinearity, based on the embedding dimension from the first step.

For a given target gene, the time-series is used to create a univariate “shadow” attractor by taking time lagged values as different coordinates in a multivariate Cartesian space (shown in Fig. 2A of the main text for example yeast genes). Although the attractors shown in the figures are all depicted in 3-dimensions, computationally we consider up to 8 lag-coordinates, i.e. an embedding dimension, *E* = 8. If the gene has deterministic dynamics, then the time-series dynamics can be predicted by nearest neighbor forecasting using simplex projection: the predicted value of a gene at the next time index is the weighted average of its E+1 nearest points on the E-dimensional univariate attractor, after following each of these points forward by one time step [47]. This is done for each time step, and each E up to E = 8. The forecast skill is measured as the Pearson correlation between predictions and observations for each E, and the optimal E is the one with the highest forecast skill. This highest forecast skill also quantifies the predictability of the dynamics. The significance of this predictability is assessed by a Wilcoxon test on the distribution of forecast errors at the optimal E having a median less than 0.

The S-map step is applied next, using the state space and univariate attractor defined in the previous step for the optimal value of embedding dimension E. For each time step, the dynamics are predicted one time step ahead using a linear model fit on all the other observations. However, the points are weighted based on their distance in the state space. When nonlinearity parameter θ = 0, all points are given equal weighting which produces the same global linear fit as a linear auto-regressive model (AR). As θ is increased, the regression becomes more locally weighted and hence more nonlinear (local weighting implies greater state dependence in the coefficients calculations). The optimal θ* is chosen to maximize forecast skill (Pearson’s correlation between the predicted values and observed values). Significance is assessed by two-sample Wilcoxon test on the forecast errors at θ* being significantly lower from the forecast errors at θ=0.

#### Concatenation and composite attractors

Empirical dynamic modeling can be improved if the reconstructed attractor is denser and this can be accomplished with more data. In the absence of obtaining a single long time series (which is not possible with synchronization), a composite attractor constructed from time series of multiple identical or similar experimental systems can be used [51]. For the yeast system, there are two realizations of wildtype dynamics, one from each batch. So that lag-coordinate vectors do not contain lags from multiple batches, composite attractors must be created with disjoint libraries. Figure S3 shows how concatenating the two wildtype time series {*Y*_WT1_, *Y*_WT2_} improves predictability and the ability to detect nonlinear dynamics. Univariate and cross map EDM analyses of wildtype dynamics reported in the main text make use of this compositing.

#### Gene-gene CCM calculations

Pairwise analysis of gene-gene associations was done with linear correlation and convergent cross mapping, CCM [14]. The basic intuition for cross mapping is described in the video clip https://www.youtube.com/watch?v=NrFdIz-D2yM. While the linear correlation is symmetric for a pair of genes *X* and *Y*, the cross map skill will generally differ in each causal direction. Calculations were performed between all genes in both yeast and mouse systems, although the main text puts particular focus on identifying genes acting on targets WHI5 (YOR083W) and YHP1 (YDR451C). As noted above, we make use of both wildtype datasets by concatenating them together, with {*Y*_WT1_, *Y*_WT2_} representing the concatenation of time series of gene *Y*. Thus, the linear correlation between genes *X* and *Y* is calculated as the Pearson’s correlation coefficient between {*X*_WT1_, *X*_WT2_} and {*Y*_WT1_, *Y*_WT2_}.

The procedure for measuring the cross map skill involves an additional step, as the embedding dimension of the predictor manifold must be chosen. First, the cross map skill from M*_X_* to *Y* is calculated for *E* from 1-8 with a prediction time of *tp* = −1 (a hind-cast cross map (see [52]), then *E\** is determined that maximizes the prediction skill measured by Pearson’s correlation between observed *Y* and predicted *Ŷ*_*tp*=−1_|**M**_***X***,***E***_. This procedure for selecting *E\** on the hind-cast skill reduces false-positives since the *E\** will be essentially random for two genes that have no dynamic relationship, but will be meaningful for two genes that are causally related.

Finally, cross mapping is performed with embedding dimension *E\** with a prediction time of *tp* = 0 (instantaneous cross map), and we measure the Pearson’s correlation (CCM ⍴) between observed *Y* and predicted *Ŷ*_*tp*=0_|**M**_***X***,***E***∗_. Note that cross mapping is reciprocal to causality [14], so cross mapping from **M***_X_* to *Y* indicates that *Y causes X*, or that the causal direction is from gene *Y* to gene *X*.

#### Co-prediction

Co-prediction measures dynamic similarity of time series by using an attractor constructed from one of the time series to predict the other. The implicit hypothesis is that the time series are different realizations of the same underlying attractor dynamics, up to a linear scaling factor. Thus, to test whether variables *X*_1_ and *X*_2_ are co-predictable one uses {*X*_1_} to predict {*X*_2_} and vice versa, normally normalizing each time series first (see Data Processing Standardization above). The skill of co-prediction, measured by the Pearson’s correlation between observed *X*_2_ to co-predicted *X*_1_, is a measure of the similarity of the dynamics of *X*_1_ and *X*_2_. This notion of using co-prediction to measure dynamic similarity has been used to distinguish between healthy and diseased heart rhythms in human infants [29].

Here, we implement co-prediction in a slightly more refined way to assess the change in dynamics between the wildtype and manipulated treatment. To assess change under the Batch 1 experiment, WT1 is taken as the control, ExM1 is taken as the manipulation, and WT2, the wildtype time series of the batch, is used as a common reference attractor. Co-prediction skill from the common reference WT2 to WT1 and ExM1 is then compared. If the dynamics of a gene are unchanged under the experimental manipulation, then WT1 and ExM1 should be equally well co-predicted from WT2. Alternatively, if the dynamics are changed under manipulation, then the WT1 should be better co-predicted than ExM1. Thus, we measure the difference in predictive skill between the co-predictions to quantify dynamic change under the experimental manipulation. The exact same procedure is flipped to assess change under batch 2 (WT2 is taken as the control, ExM2 is taken as the manipulation, and WT1 is taken as the reference attractor).

Operationally, we measure the out-of-sample predictions of *X*_WT1_ and *X*_ExM1_ from *X*_WT2_ using simplex projection [47] giving measures of prediction skill (Pearson correlation between observation and prediction) ρ_WT2→WT1_ and ρ_WT2→ExM1_, respectively. The embedding dimension for the reference attractor, is determined for each separately by choosing E* from 1 to 8 that maximizes the co-prediction skill (Pearson ρ). The difference between these, *ρ-diff* = ρ_WT2→WT1_ – ρ_WT2→ExM1_, then indicates the change in dynamics under experimental manipulation, and significance of the change in co-prediction skill is quantified by performing a two-sample Wilcoxon test on the individual forecast errors. As described above in Data Processing, note that each gene is standardized by batch so that *X*_WT1_ and *X*_WT2_ both have a range [0,1], removing batch effects, but *X*_ExM1_ and *X*_ExM2_ still reflect relative changes in level due to manipulation. Thus, this procedure covers cases where manipulation causes a change in mean level (the common focus for gene manipulation experiments) in addition to covering changes in dynamics (e.g. in peak timing or frequency) that do not result in gross changes in mean level.

**Fig. S1.**
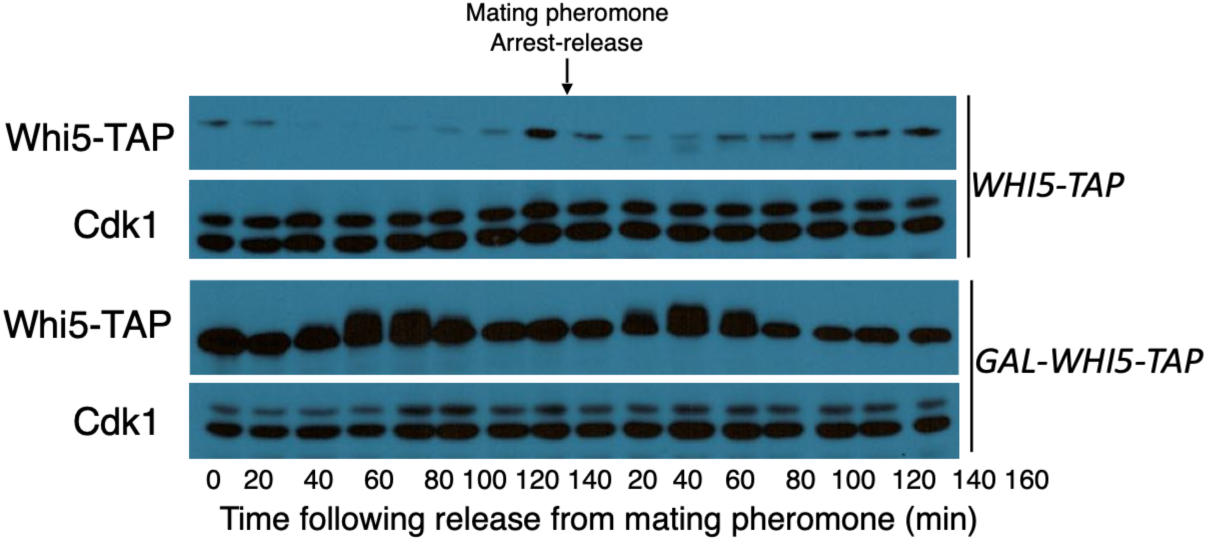
Galactose induced *WHI5* overexpression. Yeast cell cycle was synchronized by alpha factor mediated arrest. Wildtype control strain CWY 2090 BY4741 *bar1::LEU2* and a strain that overexpresses *WHI5* under the control of the GAL promoter CWY 2560 BY4741 *bar1::LEU2 whi5*::KAN were compared for the two cell cycles with resynchronization of the cell cycle with alpha factor after exit from M phase. Immunoblotting with anti *WHI5* antibody confirms overexpression of the *WHI5* protein throughout both cell cycles throughout the entire time series. Knockout of *yhp1* was confirmed by RNAseq which gave no *yhp1* reads (TPM = 0) from the *yhp1Δ*: (CWY2565*: MATa his3Δ1,leu2Δ0,met15Δ0,ura3Δ0 bar1Δ::LEU2 yhp1::KANr*) strain. Choice of genes for experimental manipulation required that the manipulation did not grossly affect the cell cycle so that it was possible to obtain time series data. As such it is important that we introduce mutations that still allow sampling of time series. For this it is important (1) that one can still synchronize the cell cycle with alpha factor and (2) that cells will still grow reasonably well and maintain synchronous growth for at least two cell cycles. WHI5 overexpression was chosen over knockout because deletion of WHI5 leads to poor growth and does not generate usable time series. Similarly, overexpression of YHP1 leads to poor synchronization of the cell cycle with alpha factor, which would also result in unusable time series.

**Fig. S2.**
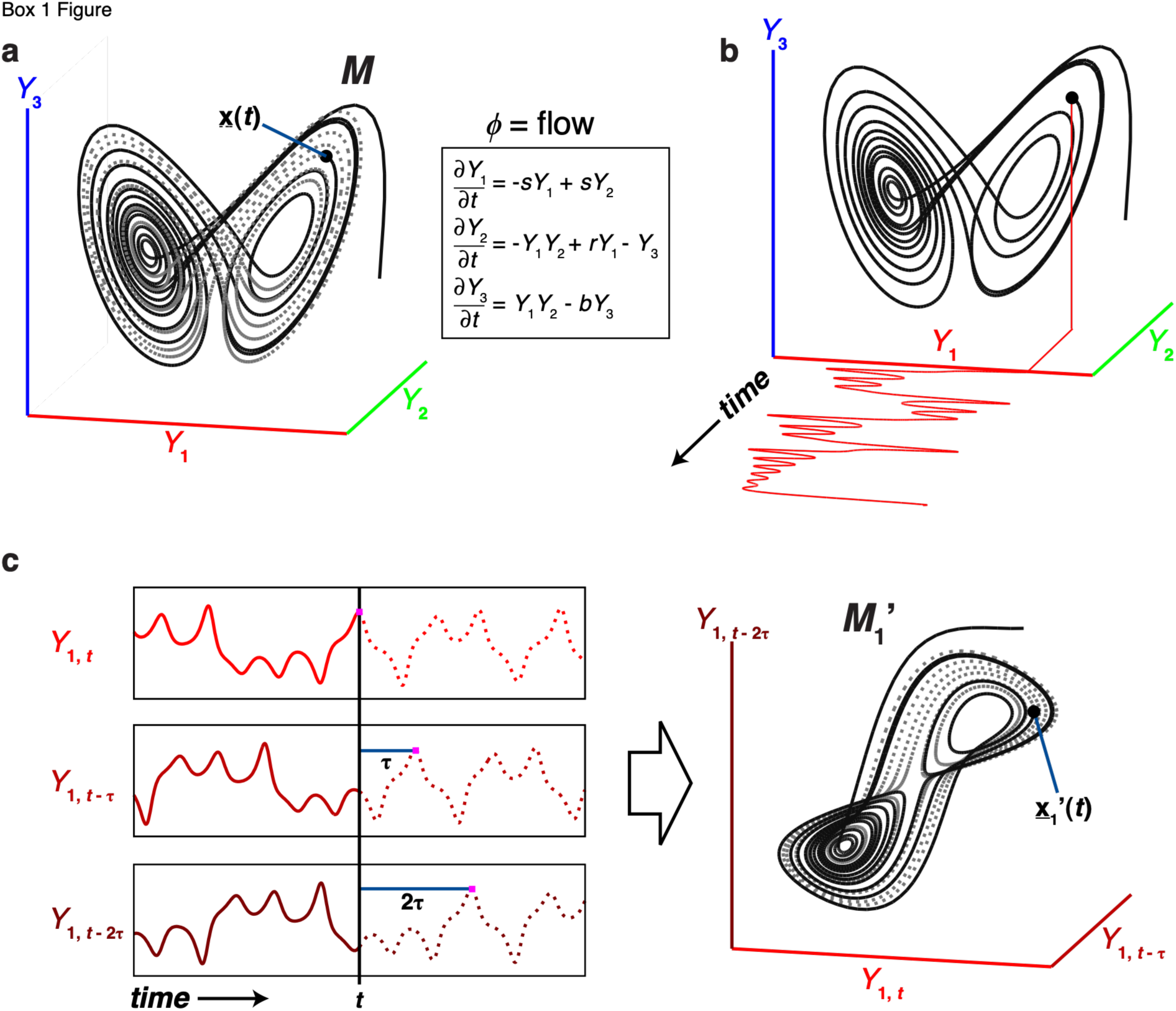
Figure legend a) Illustration of the Takens’ theorem. The Lorenz butterfly attractor example. The attractor manifold ***M*** is the set of states that the system progresses through. <u>x</u>(*t*) is the state of the system at time *t*, and the dynamics are defined by the Lorenz equations. **b)** A time series is simply a projection of the system states from ***M*** to a coordinate axis (*Y*_1_ is a state variable of the system). The manifold can be constructed from the component time series**. c)** Following Takens Theorem, lags of the time series {*Y*_1_} can act as coordinate axes to construct a shadow manifold ***M***_1_’which maps 1:1 to the original manifold ***M*** (the visual similarity between ***M***_1_’ and ***M*** is apparent). These shadow manifolds can be used for transcriptional dynamics-based prediction, identifying causal variables, taking advantage of the topological correspondence between the original system and its delay embedding through the use of correspondences of nearest neighbors in the embeddings.

**Fig. S3.**
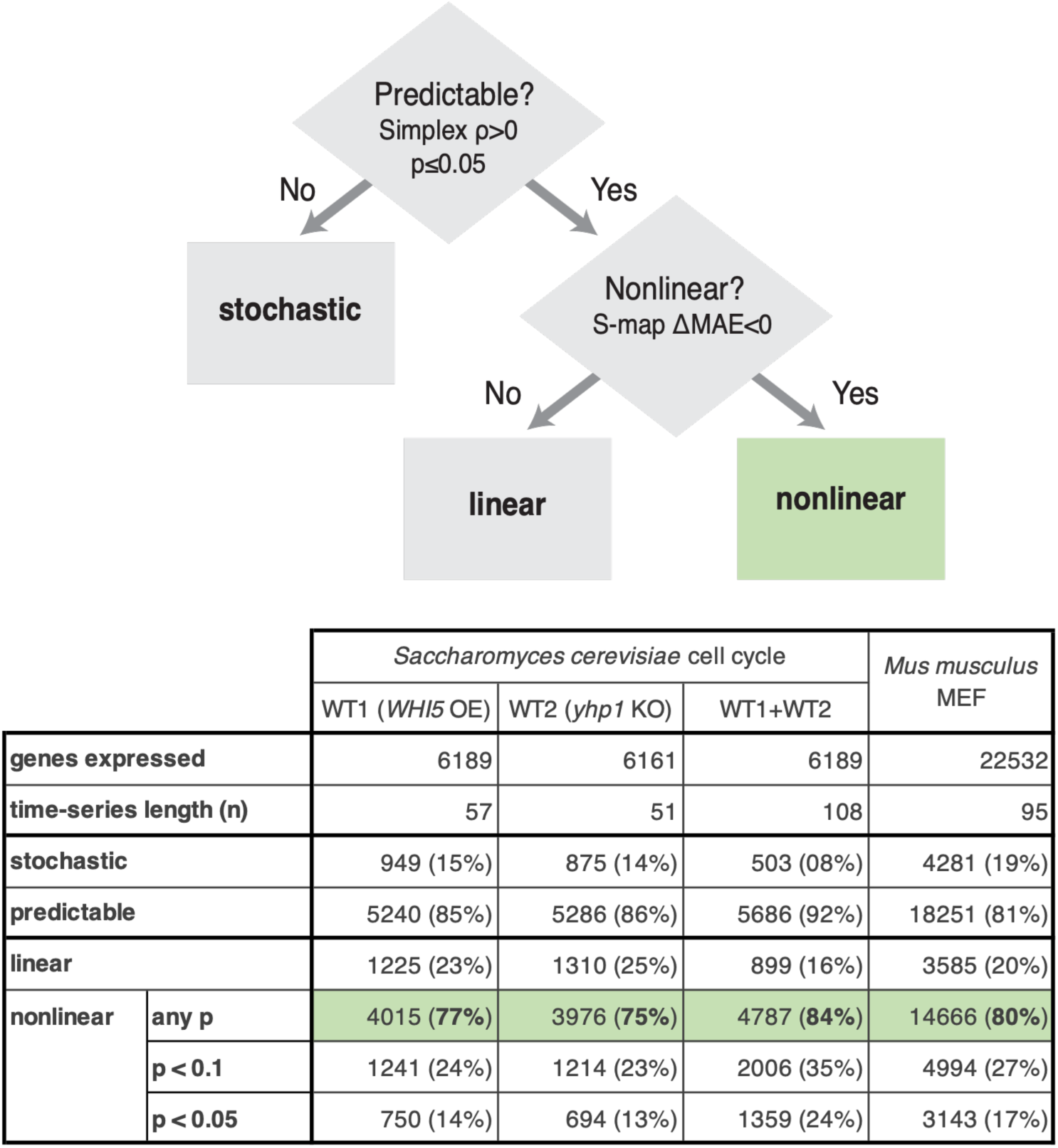
Determinism and Nonlinearity are ubiquitous in yeast and mouse gene transcription. A basic, two-step analysis evaluates predictability and nonlinearity in gene dynamics of the *S. cerevisiae* and *M. musculus* study systems. First, simplex projection is used to determine whether a gene’s expression time series is predictable (at the *p* < 0.05 level) or stochastic. Then, for predictable genes, the S-map test is applied to evaluate nonlinearity, indicated by improved prediction as models are tuned to nonlinear solutions (ΔMAE < 0). 81-92 % of genes are found to be predictable and 77-80% of predictable genes are found to be nonlinear, highlighted in green. Note that, as expected, evidence for nonlinearity in *S. cerevisiae* increases as effective time-series length is increased by concatenating the two wildtype time series.

**Fig. S4.**
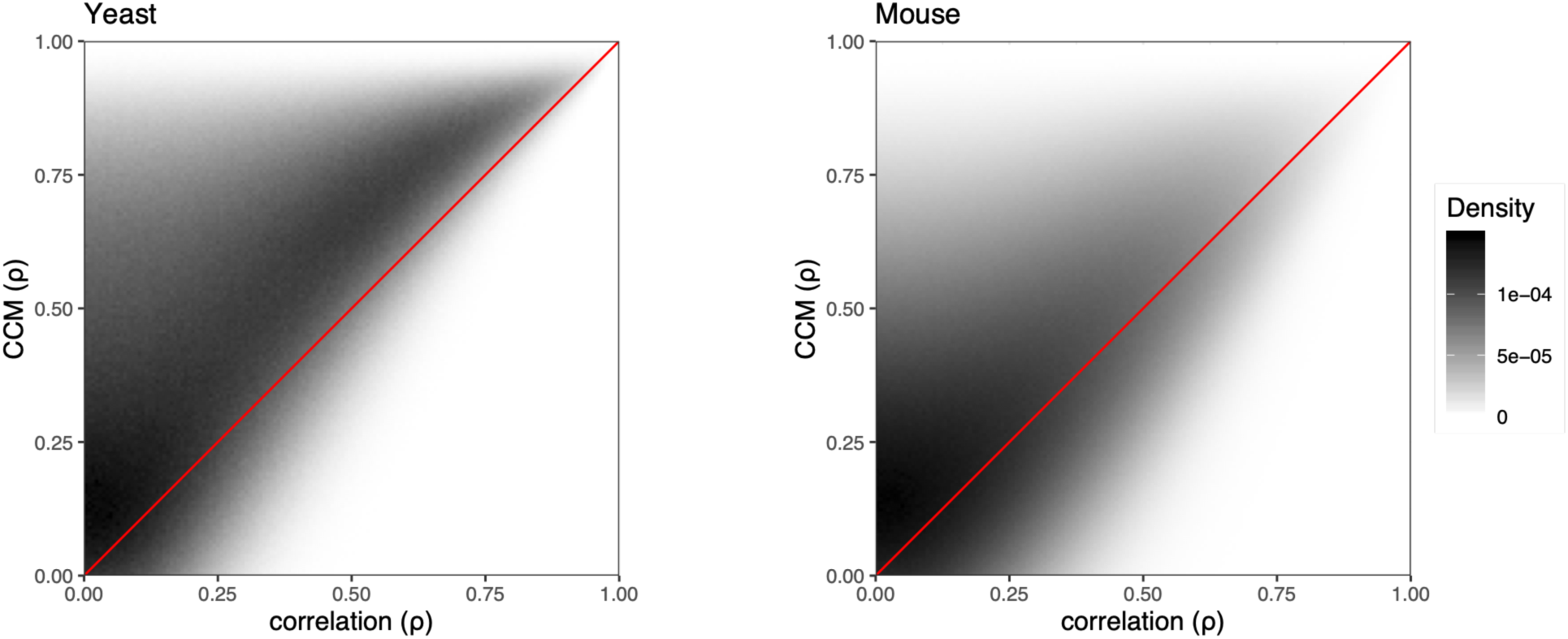
High CCM and low correlation genes are commonly found in Yeast and Mouse. Correlation and convergent cross mapping (CCM) for all pairs of expressed genes in *S. cerevisiae* (left) and *M. musculus* MEF (right) are plotted together in 2D density maps [all expressed genes or all non-linear genes?]. While in some cases CCM is comparable to correlation (density along the red 1- to-1 line), there are many cases where CCM is considerably higher than correlation, the top left quadrant. This suggests widespread occurrence causal, uncorrelated interactions invisible to static, correlation-based methods. Notice also that there are minimal highly-correlated, low-CCM pairs, indicating that CCM captures both correlated and uncorrelated interactions.

**Fig. S5.**
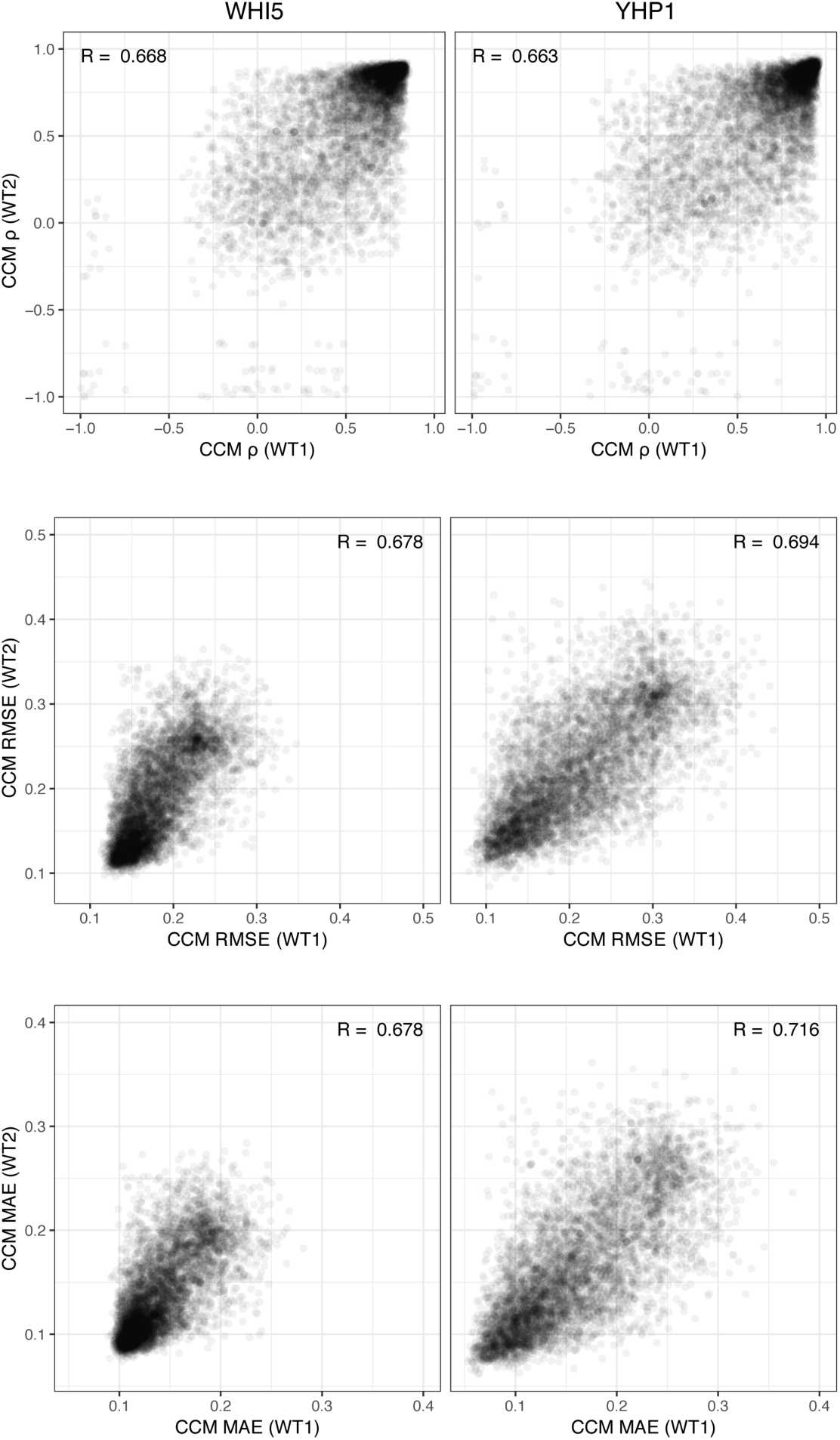
CCM consistency between *S. cervisae* experiments. Hexagonal 2D histograms show consistency between estimated CCM for targets of WHI5 (left) and YHP1 (right) in the two wildtype time series of S. cervisae. This is true whether CCM prediction is measured using Pearson’s correlation (top), root mean squared error (middle), or mean absolute error (bottom) is used to quantify CCM. Most importantly, the strong interactions (high Pearson’s ρ, low RMSE, low MAE) show the greatest consistency. For the main text, we combine the time series rather than consider the one of the experiments alone, as increased time series length produces more accurate EDM predictions (as shown univariately in Fig S1).

**Fig. S6.**
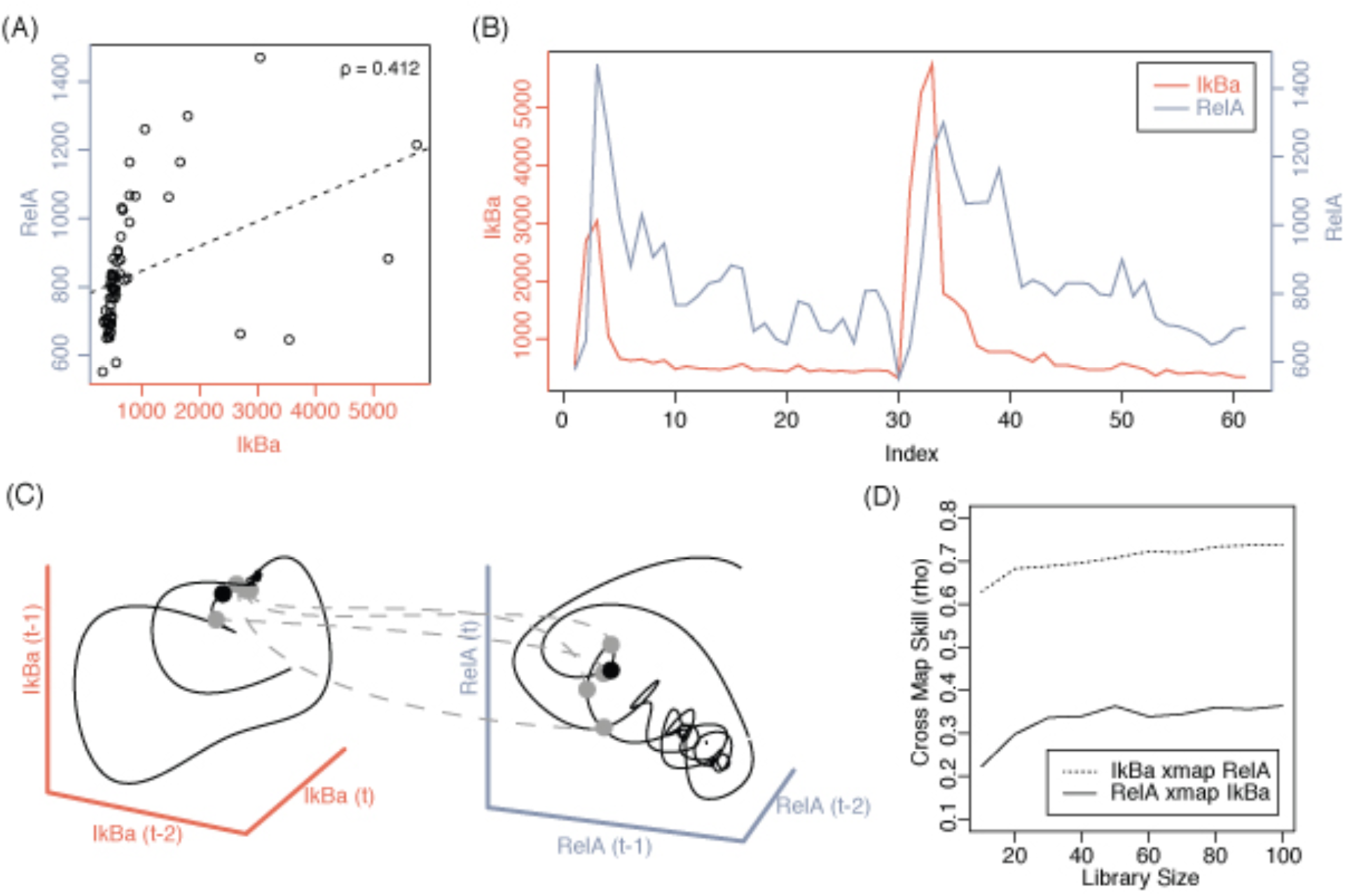
Convergent cross mapping of the mammalian NfkB system. (A) Correlation of RelA (p65) and IkBα expression values across their 2 cycle of serum response time series (B). Pearson correlation of RelA and IkBα R=0.412. (C) univariate Takens embedding in three dimensions of the shadow manifolds of RelA and IkBα using RNAseq time series data. An illustrative example of equivalent nearest neighbors of the same time index is shown as light gray dots on the manifold. The corresponding time index points are connected by dashed lines to the manifold on which they cross map. (D) Both forward and reverse regulatory relationships show convergence indicative of a convergent cross mapping signal. The causal influence of RelA onto IkBα is stronger at the transcriptional level than IkBα on RelA. This is consistent with the fact that IkBα is known to be a direct transcriptional target of the RelA/p50 heterodimer transcription complex. Although IkBα is a strong inhibitor of RelA at the posttranscriptional level where IkBα protein sequesters RelA/p50 protein and maintains it in an inactive form in the cytoplasm, IkBα was not thought affect RelA transcriptionally, which is thought to be constitutive in it’s expression. These results however hint that IkBα expression may exert a weak albeit detectable regulatory influence on RelA that may very well be indirect.

**Fig. S7.**
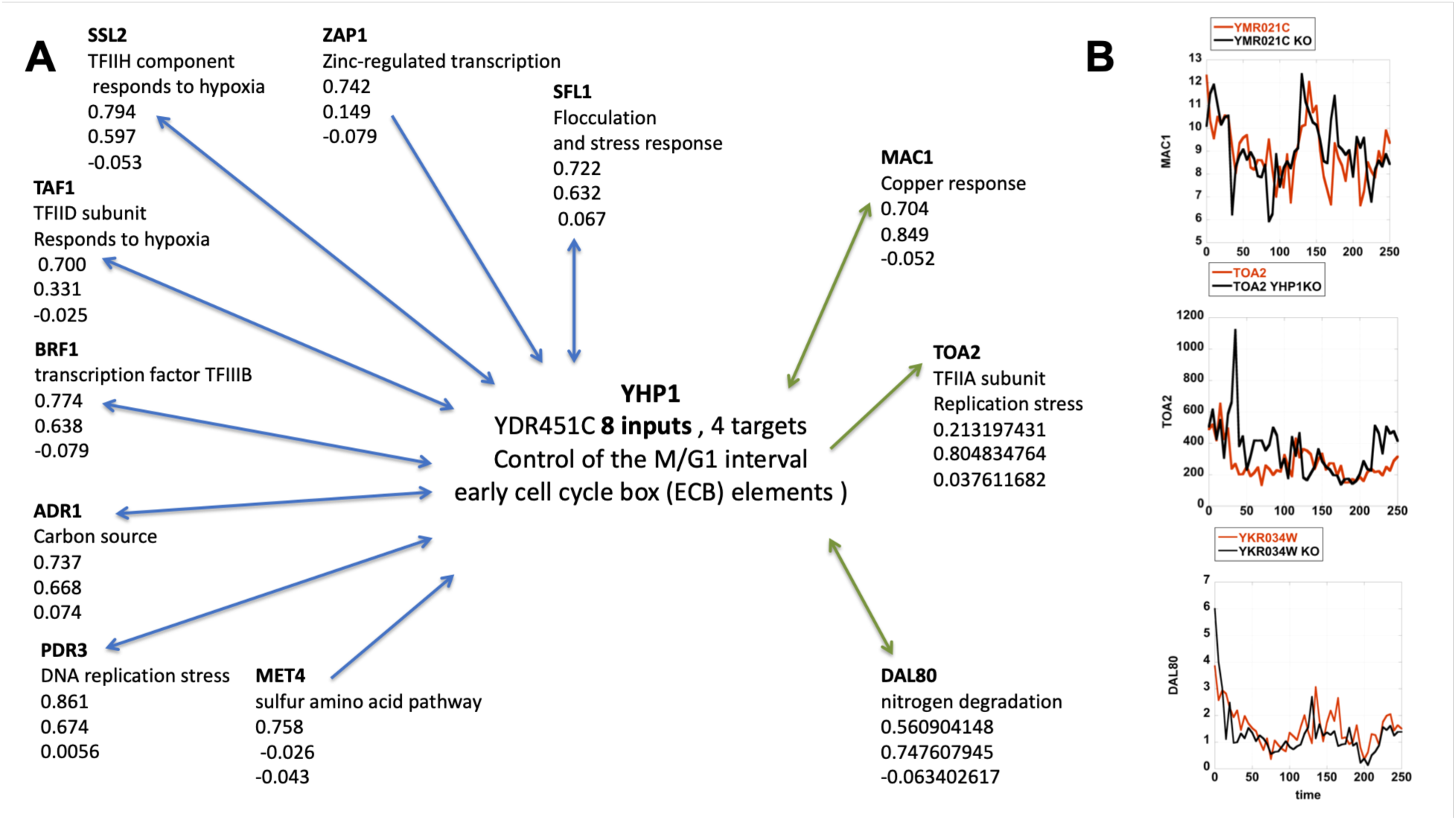
Signal integration leading to causation without correlation. (A) The CCM identified relationship of the transcription factor YHP1 with other known transcription factors. Arrows indicate the directionality of the interaction. Larger values of YHP1 cross mapping on other transcription factors indicate that the other transcription factor contains information that allows the prediction of YHP1 behavior. Hence inputs determining YHP1 behavior will in most cases have larger CCM values on the input gene. For transcriptional targets on the other hand, the behavior of the target will be better predicted by the dynamic changes of YHP1 than vice versa. As shown YHP1 integrates many physiological signals such as Zinc regulated transcription, core transcription, hypoxia, carbon source, DNA replication stress, and sulfur amino acid metabolism with the control of progression through the cell cycle. (B) Three identified uncorrelated targets all show changes in dynamics upon deletion of the YHP1 gene.

**Fig. S8.**
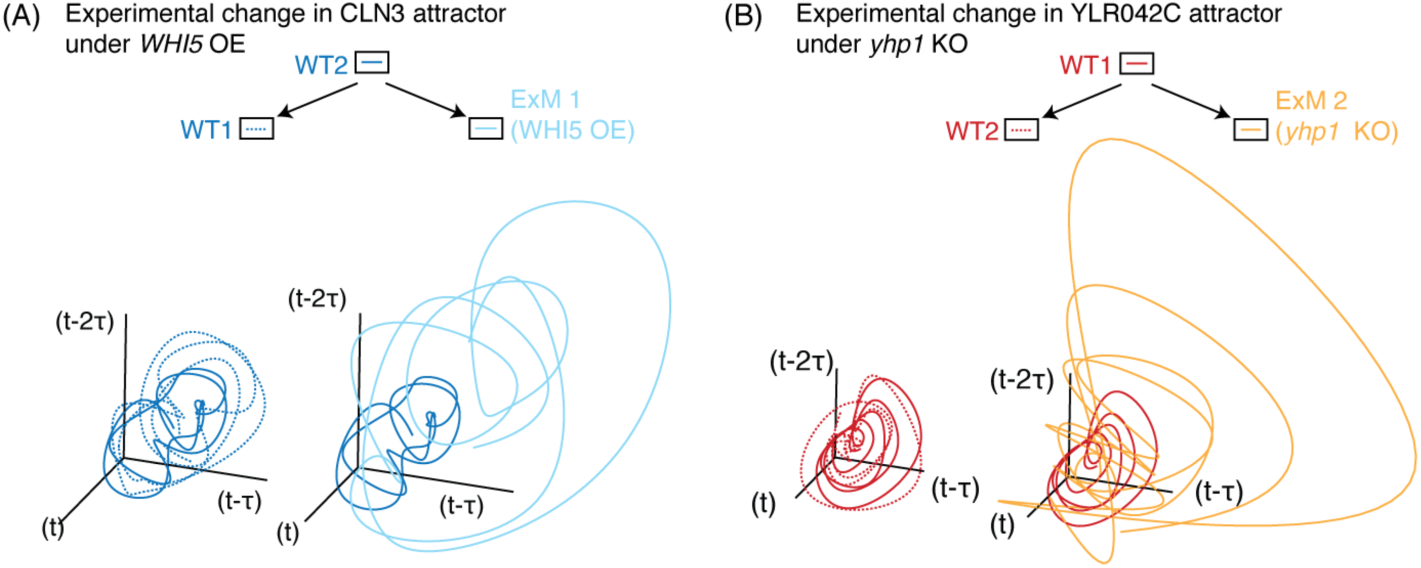
Reproduction of main text Figure 5A&B without spline smoothing of time series (all analyses in the main text use the unsmoothed raw time series as shown here) (A) The co-prediction test is graphically illustrated using the unsmoothed time series for the predicted WHI5 target, CLN3. Here the wildtype attractor (WT2) is used as the predictor (reference library) to evaluate if the manipulated attractor (ExM1) is different from the wild type attractor from that experiment (WT1). (B) Same as in (A) but where WT1 is the reference to evaluate the difference between the ExM2 and WT2 attractors for the YHP1 target YLR042C.

